# Reovirus σ1 conformational flexibility modulates the efficiency of host cell attachment

**DOI:** 10.1101/2020.06.09.143594

**Authors:** Julia R. Diller, Sean R. Halloran, Melanie Koehler, Rita dos Santos Natividade, David Alsteens, Thilo Stehle, Terence S. Dermody, Kristen M. Ogden

## Abstract

Reovirus attachment protein σ1 is a trimeric molecule containing tail, body, and head domains. During infection, σ1 engages sialylated glycans and junctional adhesion molecule-A (JAM-A), triggering uptake into the endocytic compartment, where virions are proteolytically converted to infectious subvirion particles (ISVPs). Further disassembly allows σ1 release and escape of transcriptionally active reovirus cores into the cytosol. Electron microscopy has revealed a distinct conformational change in σ1 from a compact form on virions to an extended form on ISVPs. To determine the importance of σ1 conformational mobility, we used reverse genetics to introduce cysteine mutations that can crosslink σ1 by establishing disulfide bonds between structurally adjacent sites in the tail, body, and head domains. We detected phenotypic differences among the engineered viruses. A mutant with a cysteine pair in the head domain replicates with enhanced kinetics, forms large plaques, and displays increased avidity for JAM-A relative to the parental virus, mimicking properties of ISVPs. However, unlike ISVPs, particles containing cysteine mutations that crosslink the head domain uncoat and transcribe viral positive-sense RNA with kinetics similar to the parental virus and are sensitive to ammonium chloride. Together, these data suggest that σ1 conformational flexibility modulates the efficiency of reovirus host cell attachment.

**IMPORTANCE:** Nonenveloped virus entry is an incompletely understood process. For reovirus, the functional significance of conformational rearrangements in the attachment protein, σ1, that occur during entry and particle uncoating are unknown. We engineered and characterized reoviruses containing cysteine mutations that crosslink σ1 monomers in non-reducing conditions. We found that the introduction of a cysteine pair in the receptor-binding domain of σ1 yielded a virus that replicates with faster kinetics than the parental virus and forms larger plaques. Using functional assays, we found that crosslinking the σ1 receptor-binding domain modulates reovirus attachment but not uncoating or transcription. These data suggest that σ1 conformational rearrangements mediate the efficiency of reovirus host cell attachment.

## INTRODUCTION

To infect cells, nonenveloped viruses must cross a lipid bilayer to access the cytoplasm. In most cases, the process by which nonenveloped viruses enter host cells is incompletely understood. Mammalian orthoreovirus (reovirus) rarely produces serious disease in humans, but it has been linked to the loss of oral tolerance associated with celiac disease (1), and it is being clinically evaluated for oncolytic applications. Reovirus virions are nonenveloped particles that encapsidate a segmented double-stranded RNA (dsRNA) genome within a bilayered icosahedral protein shell (2). The reovirus outer capsid is chiefly composed of μ1 and σ3 heterohexamers, while the pentameric λ2 protein spans both protein layers and anchors the trimeric σ1 protein at icosahedral vertices (3). Reovirus attachment and entry are mediated in part by σ1, which contains tail, body, and head domains (4–8) (Fig. 1). The extreme N-terminal 20-25 amino acids anchor σ1 into the viral capsid. The tail constitutes approximately one third of the molecule and forms an α-helical coiled-coil domain containing a small discontinuity (stutter) (8). The body domain is primarily formed by triple β-spiral repeats (5), while the C-terminal third of σ1 folds into a compact, eight-stranded β-barrel that forms the head domain (9).

**Figure 1.**
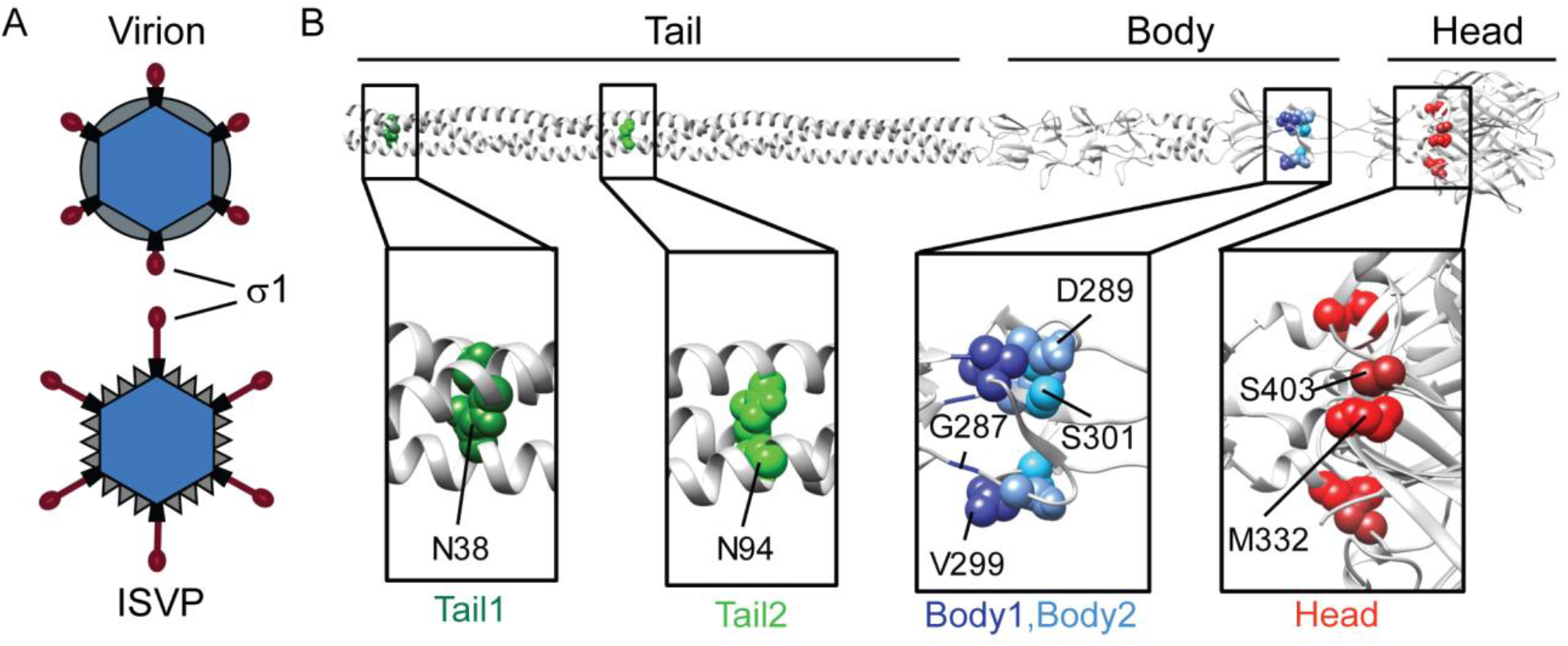
Reoviruses with engineered cysteines in the σ1 tail, body, or head domain. (A) Schematic of reovirus virion, on which σ1 adopts a compact conformation, and disassembly intermediate ISVP, on which σ1 adopts an extended conformation. (B) Full-length model of σ1 based on X-ray crystal structures of discrete σ1 domains (8). Monomers of σ1 are shown as gray ribbons, with tail, body, and head domains indicated. The locations of cysteine pairs or residues engineered in structurally adjacent sites are shown as colored spheres and labeled in magnified images below the full-length model.

To infect cells, reovirus virions adhere to sialylated glycans and the proteinaceous receptors, junctional adhesion molecule-A (JAM-A) or Nogo-66 receptor 1 (NgR1) (4, 5, 10–12). Virion binding is followed by uptake into the endocytic compartment of some types of cells via clathrin-dependent endocytosis (13–15). The σ1 head engages JAM-A within a conserved binding region, and for prototype strain type 1 Lang (T1L) σ1, the head also engages sialic acid (SA) (4, 16). For prototype strain type 3 Dearing (T3D) σ1, the body domain contains the SA-binding region (5). Reovirus attaches to cells using an adhesion-strengthening mechanism, wherein initial low-avidity σ1 interactions with SA permit lateral particle diffusion across the cell surface, which is followed by high-avidity binding to JAM-A (17). For T3D reovirus, SA binding enhances JAM-A binding capacity, possibly by triggering conformational changes in σ1 (18).

Within the endocytic compartment, reovirus virions are proteolytically converted to infectious subvirion particles (ISVPs). This conversion involves a stepwise, acid-dependent disassembly process catalyzed by cathepsin proteases (19). The first disassembly intermediate, the infectious subvirion particle (ISVP), is characterized by proteolytic removal of σ3, cleavage of μ1, and a conformational change in the σ1 protein (3, 20). Conversion from the ISVP to ISVP* involves a conformational rearrangement of the particle and μ1 autocleavage, releasing μ1 fragments that mediate formation of pores in the endosomal membrane (21–24). Additional conformational rearrangements result in σ1 release from icosahedral vertices and delivery of the transcriptionally active reovirus core into the host cell cytoplasm (21, 25, 26).

Evidence for a change in σ1 conformation comes from comparing negative-stain electron microscopy (EM) images and cryo-EM reconstructions of virions and ISVPs (3, 20). Filamentous structures frequently protrude from ISVPs but not from virions in negative-stain EM images, and electron density extending radially from icosahedral vertices is more elongated in ISVPs and more knob-like in virions in cryo-EM image reconstructions. These observations suggest that σ1 exists in a more compact form on virions and an extended form on ISVPs, leading to the hypothesis that σ1 extension in the endosome contributes to viral escape into the cytosol. However, the functional consequences of σ1 conformational rearrangements that may be triggered during attachment and virion-to-ISVP conversion are unknown. To identify contributions of conformational rearrangement to reovirus replication processes, we engineered, recovered, and characterized recombinant reoviruses containing cysteine mutations at sites in the tail, body, and head domains that formed crosslinking disulfide bridges between adjacent monomers within the trimer. Our results suggest that stabilizing σ1 alters reovirus behavior with respect to the key replication process of receptor binding.

## RESULTS

### Reovirus disulfide mutants encapsidate crosslinked σ1

To determine the functional importance of σ1 conformational mobility, we introduced cysteine mutations into structurally adjacent sites in the σ1 tail, body, and head domains using plasmid-based reverse genetics (Fig. 1B). Cysteine mutations in the tail were engineered at residues that coordinate chloride ions in the α-helical coiled-coil interior (8) and were anticipated to result in the formation of disulfide bridges yielding crosslinked σ1 dimers in oxidizing environments. Cysteine pairs introduced into the body and head domains were anticipated to crosslink σ1 in a trimeric conformation. All cysteine mutant viruses were successfully recovered and replicated to reasonable titers following two passages in murine L929 fibroblasts (L cells) (Table 1).

**TABLE 1.** Reovirus titer following two L-cell passages.

| Virus name | $\sigma 1$ mutations | Clone | Titer (pfu/ml) |
| --- | --- | --- | --- |
| rsT1L | | 1 | $5.2 \times 10^7$ |
| | | 2 | $5.6 \times 10^7$ |
| Tail1 | N38C | 1 | $5.4 \times 10^7$ |
| | | 2 | $5.9 \times 10^7$ |
| Tail2 | N94C | 1 | $5.3 \times 10^8$ |
| | | 2 | $3.3 \times 10^8$ |
| Body1 | G287C, V299C | 1 | $1.3 \times 10^6$ |
| | | 2 | $6.1 \times 10^5$ |
| Body2 | D289C, S301C | 1 | $2.8 \times 10^6$ |
| | | 2 | $9.7 \times 10^5$ |
| Head | M332C, S403C | 1 | $5.9 \times 10^8$ |
| | | 2 | $2.7 \times 10^7$ |
| HeadΔSA | M332C, S370P, | 1 | $5.7 \times 10^7$ |
| | Q371E, S403C | 2 | $2.7 \times 10^8$ |

To characterize σ1 crosslinking and encapsidation phenotypes, we purified particles of the cysteine mutant viruses. We easily purified all viruses except the Head mutant (M332C, S403C), which sometimes yielded faint bands following cesium-chloride gradient centrifugation (Fig. 2A) that did not appear to contain viral proteins when analyzed by immunoblotting. We hypothesized that the engineered cysteine mutations crosslink the T1L σ1 head domain, which contains a glycan-binding site, thus altering virus glycan-binding properties. Accordingly, incubation of the infected L-cell pellet with neuraminidase (NA) prior to reovirus extraction and gradient ultracentrifugation resulted in a banding pattern that was indistinguishable from that of parental rsT1L, suggesting that glycan binding prevents Head mutant release from a lipid-containing fraction of the cell pellet.

**Figure 2.**
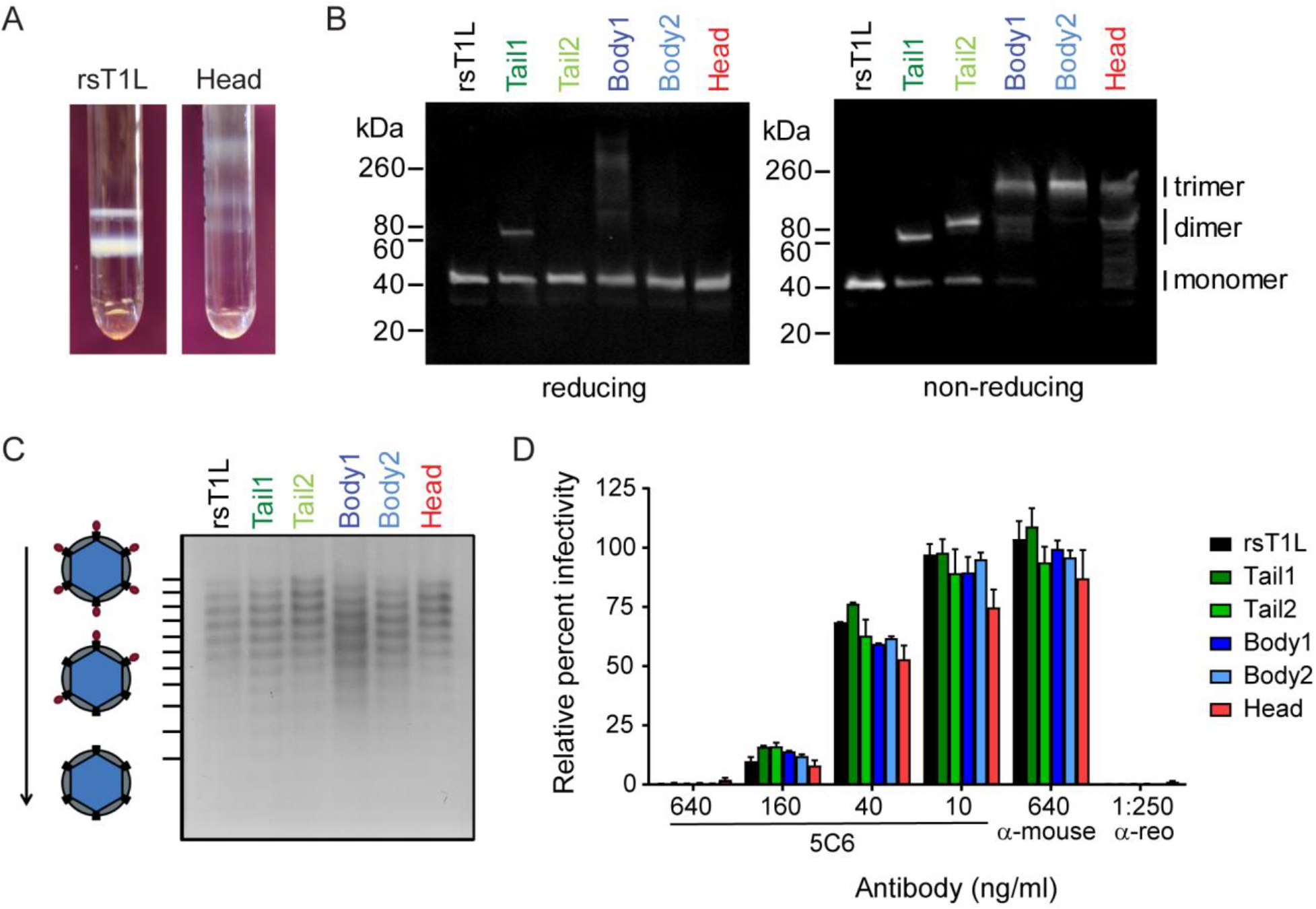
Cysteine mutant virus characterization. (A) Banding pattern for rsT1L or the Head mutant following standard reovirus purification by cell lysis, organic solvent extraction, and cesium-chloride gradient ultracentrifugation. (B) Cysteine mutations were designed to allow disulfide bond formation between σ1 monomers in the trimer. Disulfide bond formation was confirmed by boiling equivalent particle numbers of parental or cysteine mutant viruses in the presence (reducing) or absence (non-reducing) of β-mercaptoethanol, followed by SDS-PAGE and immunoblotting using antibodies specific for the σ1 head domain and fluorescently labeled secondary antibodies. Molecular weight markers and the approximate location of monomeric, dimeric, and trimeric σ1 species are indicated. (C) Equivalent numbers of purified viral particles were resolved by agarose gel electrophoresis and stained with colloidal Coomassie. Particles migrate as distinct bands based on the number of encapsidated σ1 molecules. (D) Equivalently infectious concentrations of purified viral particles were incubated with increasing concentrations of mAb 5C6, anti-mouse IgG as a negative control, or polyclonal reovirus antiserum (α-reo) as a positive control, prior to adsorption on L-cell monolayers. 5C6 is a conformation-specific mAb that binds the σ1 head domain across the interface between two monomers, thereby preventing cell attachment. Cells were fixed at ∼ 18 h postadsorption and scored for DAPI and reovirus antigen by indirect immunofluorescence. Results are expressed as mean percentage of virus-infected cells, relative to mock-treated cells, from four fields of view per well in duplicate wells. Error bars indicate standard deviation.

To determine whether the σ1 proteins encapsidated by the cysteine mutant viruses displayed the anticipated crosslinking phenotypes, we analyzed purified reovirus particles in the presence or absence of the reducing agent β-mercaptoethanol by immunoblotting with an antiserum specific for the σ1 head domain. As anticipated, cysteine mutations in the tail yielded crosslinked σ1 dimeric or monomeric species in oxidizing environments, while cysteine pairs introduced into the body and head domains yielded trimeric or sometimes dimeric σ1 species (Fig. 2B). In a reducing environment, σ1 mutants were present solely as monomers, except for Tail1 (N38C) and Body1 (G287C, V299C), which retained a minimal level of multimeric character. To determine whether cysteine mutant viruses encapsidated equivalent numbers of σ1 molecules, we resolved purified particles using agarose gel electrophoresis. All mutants encapsidated numbers of σ1 trimers comparable to parental rsT1L, typically 6-12 (Fig. 2C). To evaluate folding of σ1 trimers, we employed a microneutralization assay using conformation-specific monoclonal antibody 5C6, which binds across the interface between two σ1 monomers (27). Infection of L cells by parental rsT1L and all cysteine mutant viruses was efficiently neutralized by 5C6 but not by control mouse IgG, suggesting encapsidated σ1 forms properly folded trimers on virus particles (Fig. 2D). Together, these observations suggest the cysteine mutant viruses encapsidate equivalent numbers of folded σ1 trimers but differ in σ1 crosslinking phenotypes under oxidizing conditions.

### Some reovirus disulfide mutants display enhanced replication kinetics and form large plaques

To compare replication kinetics, we adsorbed L cells with 0.1 PFU/cell of rsT1L or each of the cysteine mutants and quantified virus titers during a time course. Titer decreased through 8 h post adsorption, began to increase at 12 h, and continued to increase through 24 h for all viruses (Fig. 3A). Although starting titers were nearly identical, titers were significantly increased for Tail1 at 20 h, for Body2 (D289C, S301C) at 20 and 24 h, and for the Head mutant at every time point except 16 h post-adsorption relative to rsT1L (*P* < 0.05 by unpaired *t* test). While determining virus titers, substantial differences in plaque size were evident (Fig. 3B). Whereas all rsT1L plaques were small, Tail1, Body2, and Head mutants had variable plaque sizes, with most plaques small for Body2 and most plaques large for Head (Fig. 3C). These differences in diameter were statistically significant for Tail1 and Head but not Body2. Inclusion of proteases, like chymotrypsin, in the plaque assay overlay yields larger plaques in fewer days, presumably by converting progeny virions to ISVPs upon release (28, 29). To determine whether conversion of virions to ISVPs could alleviate differences in plaque size, we quantified plaque diameter after including chymotrypsin in the overlay. We found that plaque size differences among the viruses largely disappeared under these conditions, although plaque diameter remained significantly larger for the Head mutant compared with rsT1L (Fig. 3D). Together, these findings suggest that engineered cysteines that crosslink the tail, body, and head domains of σ1 in oxidizing environments are not disadvantageous and can, in some cases, confer an advantage in viral replication and spread in cultured cells.

**Figure 3.**
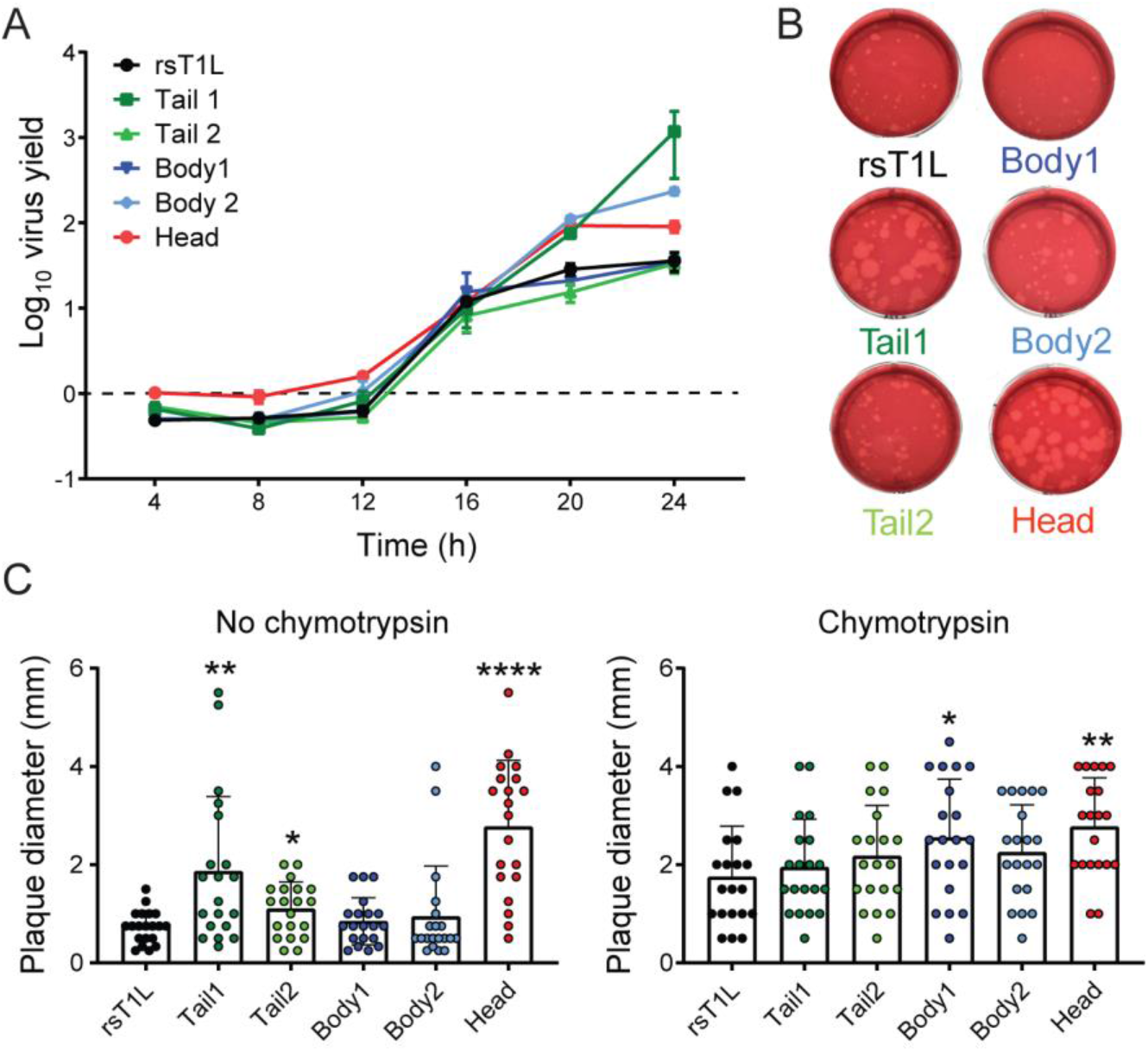
Cysteine mutant virus replication and spread. (A) L929 cells were adsorbed with rsT1L or cysteine mutant viruses at an MOI of 0.1 PFU/cell. After adsorption, unbound virus was removed, fresh medium was added, and cells were incubated at 37°C for the times shown prior to lysis by two rounds of freezing and thawing. Viral titers in cell lysates were determined by plaque assay. Virus yield was determined by dividing the average virus titer at a given time point by that at 0 h. Error bars indicate standard deviation. (B) Images of individual wells from a plaque assay. (C-D) For each virus strain, the diameters of 20 plaques were measured in mm following overlay with an agar medium mixture in the absence (C) or presence (D) of chymotrypsin, which promotes conversion of virions to ISVPs. Bars show mean and standard deviation, and dots indicate individual measurements. Values that differ significantly from rsT1L by unpaired *t* test are indicated by * (*P* < 0.05), ** (*P* < 0.01), or **** (*P* < 0.0001).

### The Head mutant large-plaque phenotype is independent of sialic acid-binding capacity

Inconsistency in purification of the Head mutant in the absence of NA (Fig. 2A) suggested that this virus may have altered SA-binding properties. Since the T1L SA-binding site resides in the σ1 head domain (4), crosslinking head monomers may change virion SA-binding properties. To distinguish between phenotypes mediated by altered SA-binding properties and crosslinking of the head domain, we engineered a Head mutant reovirus containing point mutations (S370P, Q371E) that decrease SA binding efficiency (4). HeadΔSA mutant particles encapsidate similar numbers of σ1 molecules as rsT1L and the Head mutant (Fig. 4A), and HeadΔSA σ1 trimers are crosslinked in non-reducing but not reducing conditions (Fig. 4B). However, the HeadΔSA mutant has reduced capacity to agglutinate human red blood cells compared with the Head mutant, suggesting it has reduced SA-binding capacity (Fig. 4C). Indeed, approximately eight times as many HeadΔSA as Head mutant particles are required to agglutinate red blood cells (Table 2). Surprisingly, the Head mutant appears to have reduced SA-binding capacity compared with rsT1L (Figs. 2A, 4C). Approximately four times as many Head particles as rsT1L particles are required to agglutinate red blood cells (Table 2). While reduced SA-binding capacity would fail to explain difficulties in purification of the Head mutant in the absence of NA, it is possible that purification in the presence of NA yielded particles in which some SA binding sites were occupied by glycan prior to conducting hemagglutination assays. In either case, these collective observations suggest that HeadΔSA virions resemble Head virions, except that they have reduced SA-binding capacity.

**Figure 4.**
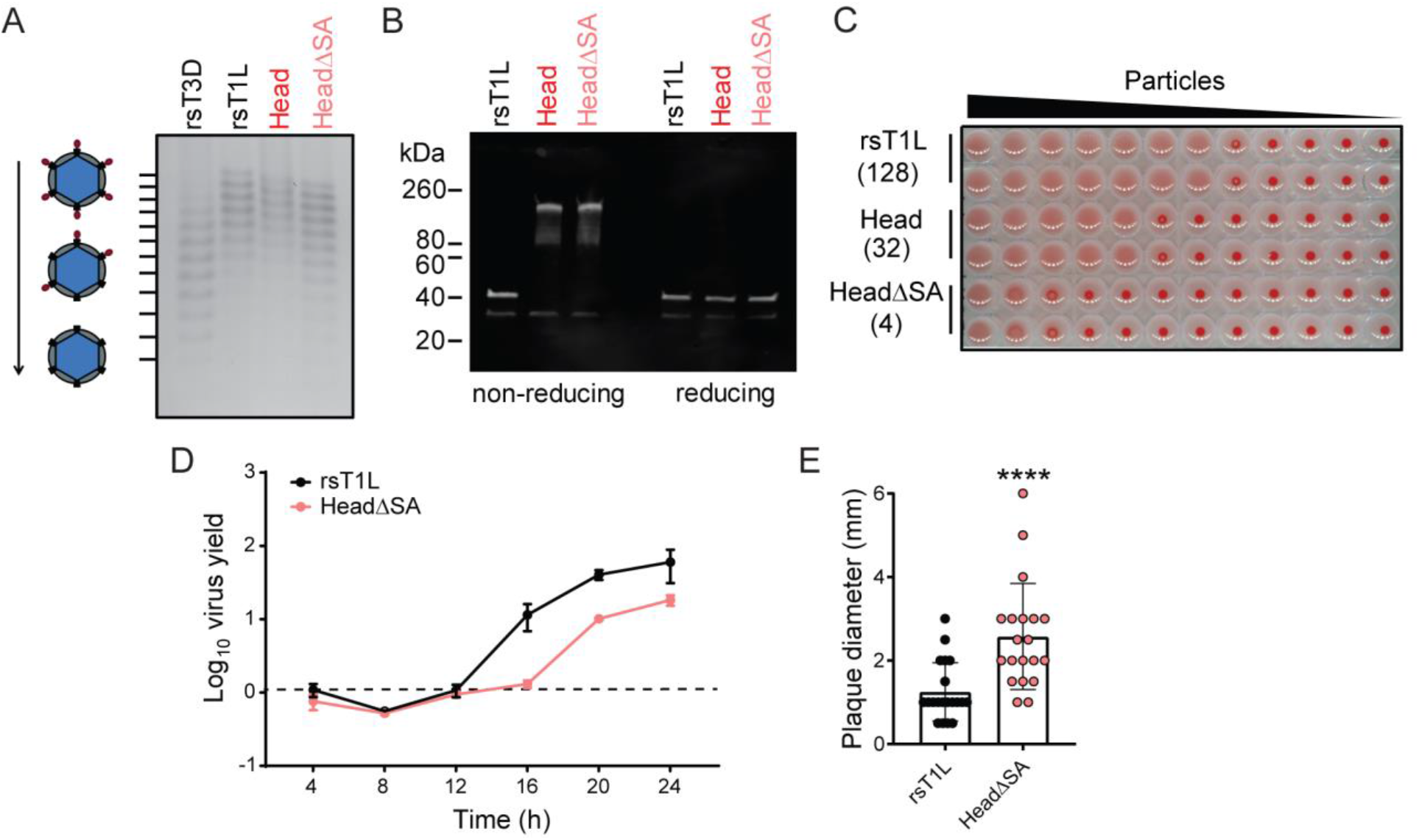
HeadΔSA mutant characterization. (A) Equal numbers of purified viral particles were resolved by agarose gel electrophoresis and stained with colloidal Coomassie. Particles migrate as distinct bands based on the number of encapsidated σ1 molecules. (B) Disulfide bond formation was confirmed by boiling equivalent particle numbers of parental or cysteine mutant viruses in the absence (non-reducing) or presence (reducing) of β-mercaptoethanol, followed by SDS-PAGE and immunoblotting using antibodies specific for the σ1 head domain and fluorescently labeled secondary antibodies. Molecular weight markers are indicated. (C) Equal numbers of purified reovirus particles were serially diluted, incubated with a 1% solution of human erythrocytes at 4°C for 4 h, and scored for HA in duplicate. Erythrocyte shields indicate HA, and erythrocyte buttons indicate absence of erythrocyte cross-linking. HA titer is shown in parentheses. HA titer is defined as 2.5 × 10^10^ particles divided by the number of particles per HA unit; one HA unit equals the minimum particle number sufficient to produce HA. (D) L929 cells were adsorbed with rsT1L or HeadΔSA at an MOI of 0.1 PFU/cell. After adsorption, unbound virus was removed, fresh medium was added, and cells were incubated at 37°C for the times shown prior to lysis by two rounds of freezing and thawing. Viral titers in cell lysates were determined by plaque assay. Virus yield was determined by dividing the average virus titer at a given time point by that at 0 h. Error bars indicate standard deviation. (E) Diameters of 20 plaques were measured in mm following overlay with an agar medium mixture. Bars show mean and standard deviation, and dots indicate individual measurements. HeadΔSA plaques are significantly larger than those of rsT1L by unpaired *t* test. ****, (*P* < 0.0001).

**TABLE 2.**
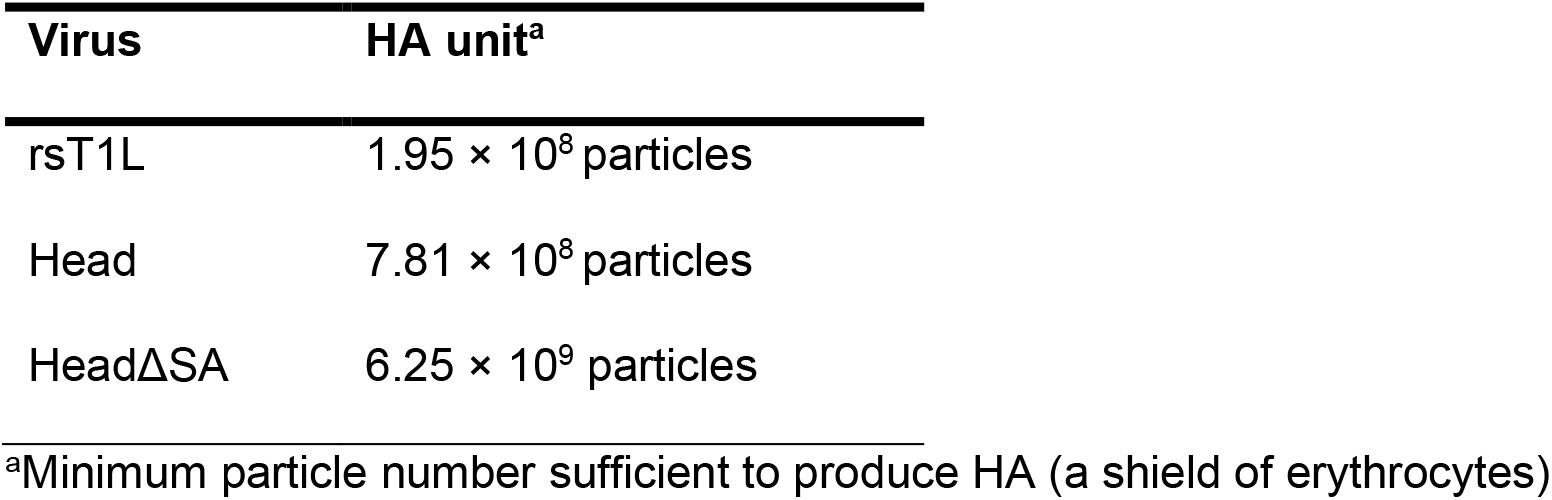
Viral hemagglutination titer using human red blood cells.

To compare replication kinetics, we adsorbed L cells with 0.1 PFU/cell of rsT1L or HeadΔSA and quantified virus titer during a time course. Virus yields decreased through 8 h post-adsorption and began to increase at 12 h for both viruses (Fig. 4D). Unlike the head mutant, HeadΔSA exhibited a delay in replication relative to rsT1L, with significantly reduced yields at 20 h post-adsorption (*P* < 0.01 by unpaired *t* test). However, like the plaques of the Head mutant, HeadΔSA plaques were significantly larger than those of rsT1L (Fig. 4E). These findings indicate that efficient SA binding is dispensable for mediating the enhanced-spread phenotype of the Head mutant in plaque assay conditions but may be required to facilitate enhanced replication.

### Mutant phenotypes do not result from particle aggregation

We thought it possible that the enhanced replication and large-plaque phenotypes of the Tail1, Body2, and Head cysteine mutants could result from particle aggregation, which could enhance replication and spread via concurrent target cell infection with multiple particles. To exclude this possibility, we compared particles of rsT1L and the cysteine mutants using dynamic light scattering (DLS). For every virus preparation, we detected one major peak with the expected hydrodynamic radius, and there was no evidence of aggregation (Fig. 5A and Table 3) (30, 31). In all cases, a smaller peak also was detected. For Tail2 (N94C) and Body2, the intensity of this peak was sufficient to define one or more replicates as polydispersed (Table 3). Effects of this significant smaller peak are reflected in reduced autocorrelation with rsT1L (Fig. 5B). The predicted molecular weight of peak 1 is consistent with that of outer-capsid protein σ3 and may represent partial particle uncoating within the sample preparation. Excellent autocorrelation with rsT1L was observed for Tail1, Body1, Head, and HeadΔSA mutants (Fig. 5B). These observations provide evidence that aggregation is not responsible for the enhanced replication kinetics and large-plaque phenotypes of the Tail1, Body2, or Head cysteine mutants and suggest potential differences in uncoating for the Tail2 and Body2 mutants.

**Figure 5.**
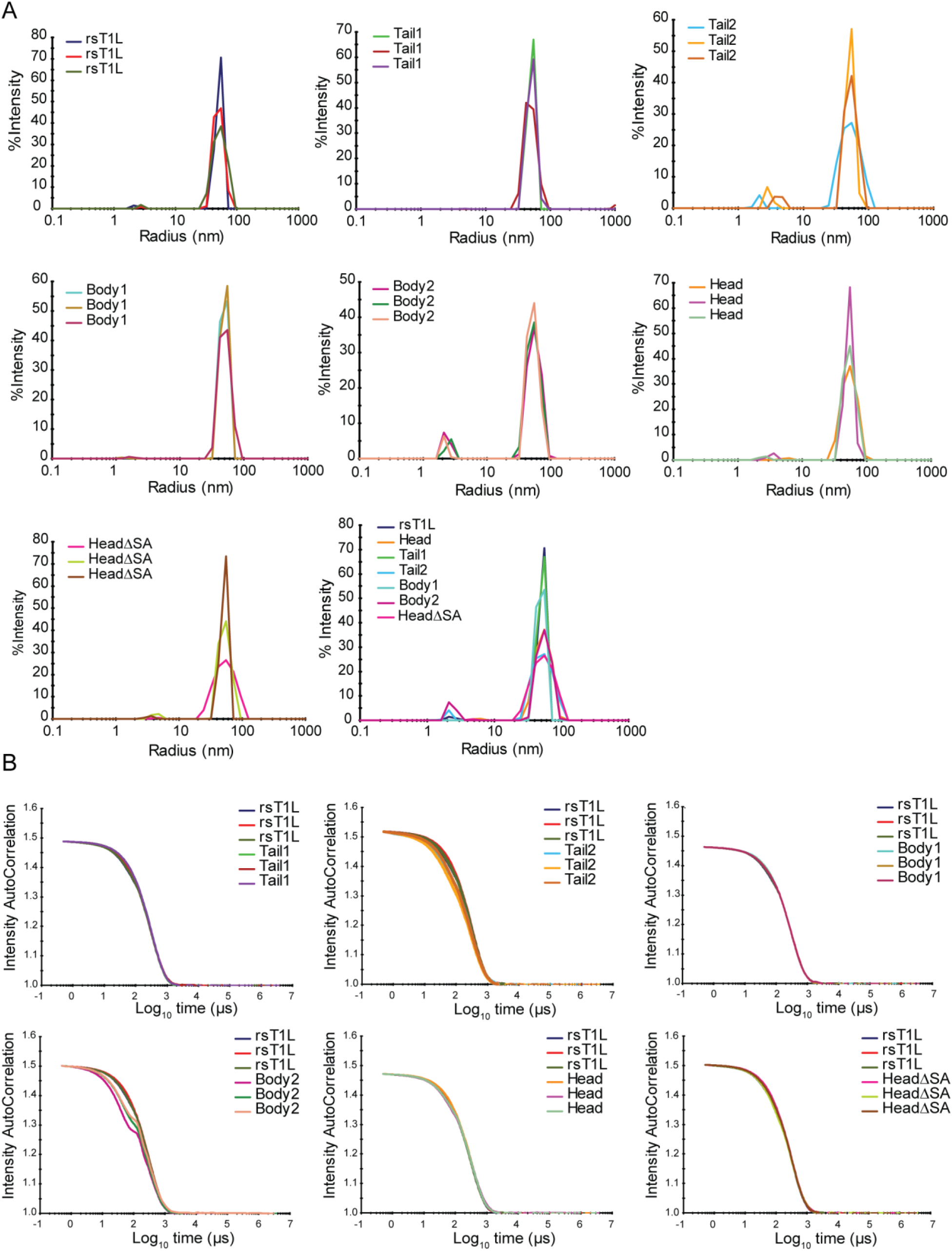
Dynamic light scattering data for rsT1L and cysteine mutants. (A) Particle size distribution curves expressed as percentage total light scattering intensity. rsT1L or cysteine mutants were analyzed by DLS in three replicate experiments. The size distribution profiles from the three replicates for a specific virus or a single representative replicate for all viruses are shown overlaid. Average hydrodynamic radii are shown in Table 3. (B) Autocorrelation curves for rsT1L overlaid on those for each cysteine mutant virus from three replicate experiments.

**TABLE 3.**
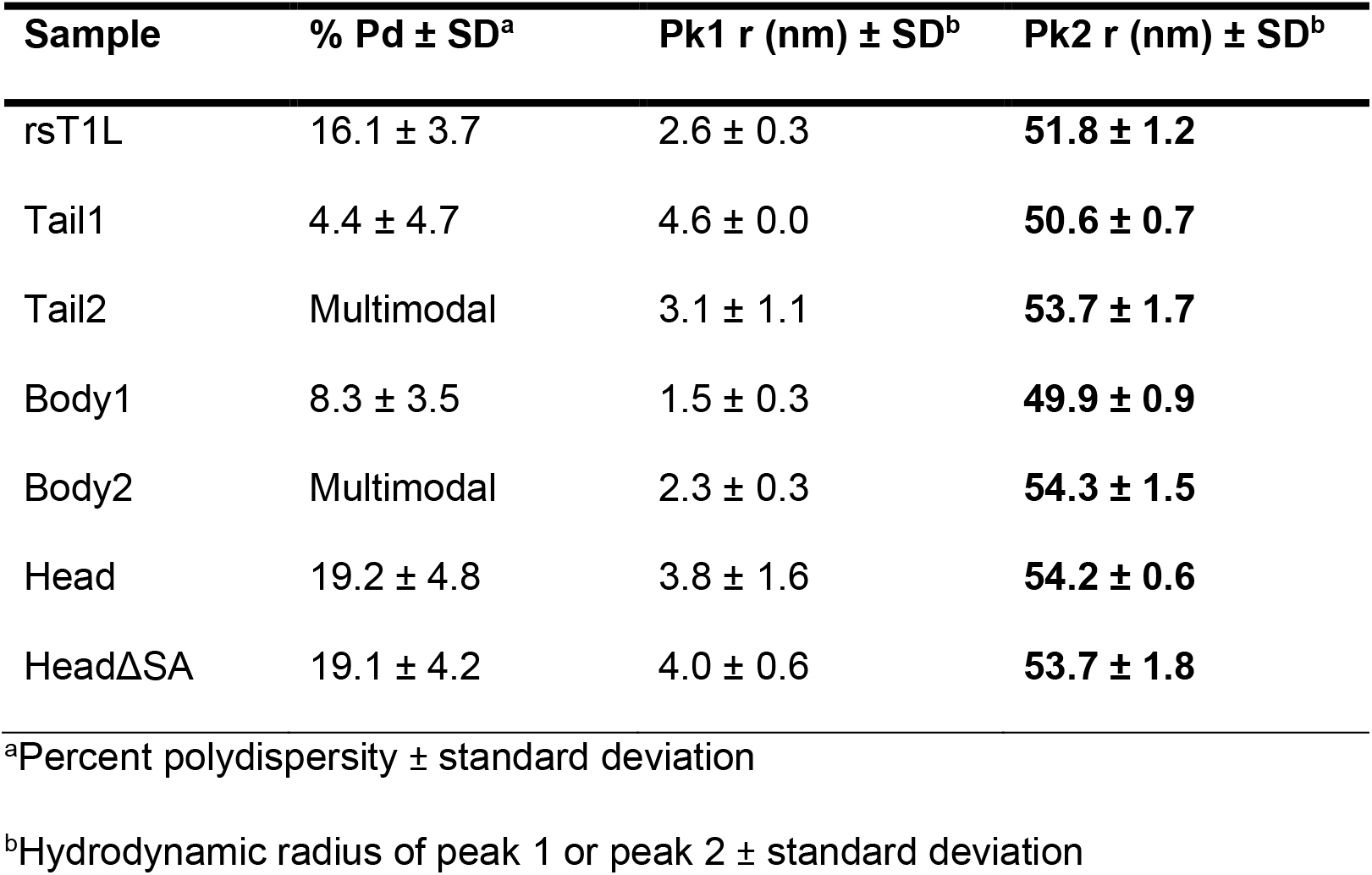
Dynamic light scattering statistics.

### The Head mutant binds JAM-A with higher avidity than parental rsT1L

The enhanced replication and large-plaque phenotypes observed for some σ1 cysteine mutants are reminiscent of ISVPs. ISVPs with a serotype 3 σ1 have higher avidity for JAM-A than do virions, which is hypothesized to result from the conformational change in σ1 that accompanies virion-to-ISVP conversion (18). We hypothesized that serotype 1 ISVPs also may have a higher JAM-A avidity than virions and that our crosslinked Head mutant adopts a conformation like that of serotype 3 ISVPs. To quantify JAM-A avidity, we used single-virus force spectroscopy, which is based on atomic force microscopy (AFM). We focused our analyses on the Head mutant, as it consistently formed large plaques, exhibited enhanced replication kinetics, and had excellent autocorrelation with rsT1L by DLS (Figs. 3-4). Single virion particles (rsT1L, Head, or HeadΔSA) were covalently attached to the AFM tip, and interactions with a surface coated with purified JAM-A were kinetically and thermodynamically probed (Fig. 6A). The AFM tip was cyclically projected toward and retracted from the JAM-A coated surface. The force acting between the functionalized tip and the surface (expressed in piconewtons) was monitored over time, resulting in force vs. time curves (Fig. 6B) (32, 33). Upon retraction, binding events were frequently observed, and their magnitudes were extracted giving the binding force observed between virions and JAM-A. By comparing the three types of virions, we observed a binding probability as follows: HeadΔSA ≈ Head >> rsT1L (Fig. 6C), confirming that crosslinking the head domain influences JAM-A binding. Next, we studied the kinetic properties of the established bonds by probing the interaction at various loading rates (i.e., the force applied over time). Using the Bell-Evans model (34), we fit the data and extracted the dissociation rate (k_off_) and the distance to the transition state (x_u_) for the three virion types, rsT1L (Fig. 6D), Head (Fig. 6E) and HeadΔSA (Fig. 6F). Overall, we observed a decrease in the distance to transition state upon crosslinking of σ1 monomers, indicating a reduction of the conformational variability following binding to JAM-A as well as a reduction of the dissociation rate, suggesting more stable interactions. Finally, by monitoring the influence of contact time on binding probability (Fig. 6G-I), we observed an increase in the association rate (k_on_) for both mutants, suggesting that crosslinking the head domain facilitates JAM-A binding, as also highlighted by a significant decrease of the equilibrium dissociation constant. Therefore, a single-virus biophysical approach indicates that crosslinking the σ1 head enhances the avidity of reovirus T1L for JAM-A.

**Figure 6.**
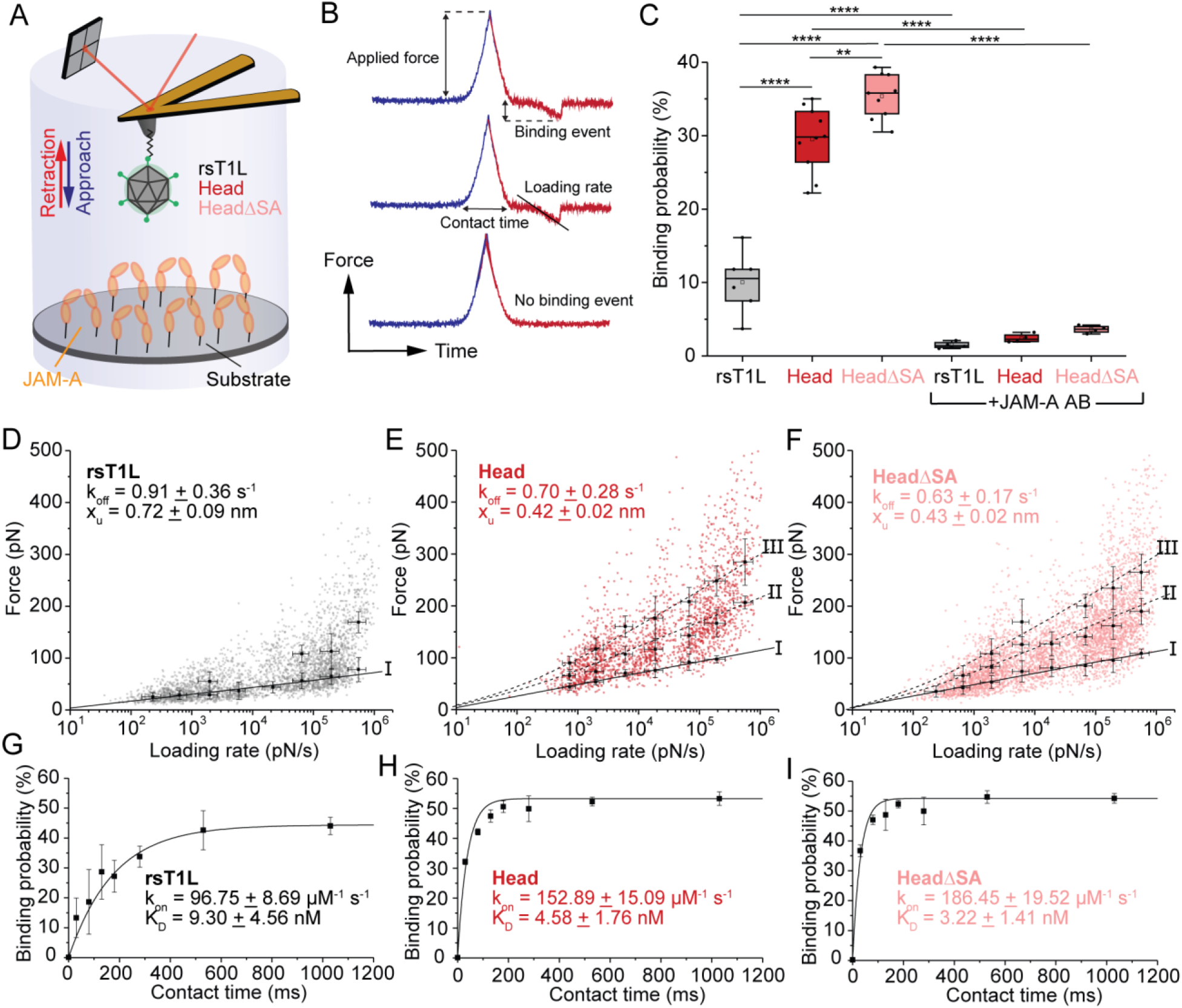
Head mutant avidity for JAM-A quantified by AFM. (A) Principle of force distance curve-based atomic force microscopy (FD-based AFM). An AFM tip functionalized with a PEG spacer fused to the virus of interest is projected toward and retracted from a surface coated with JAM-A. (B) Force-time curve from which the loading rate (LR) can be extracted from the slope of the curve just before bond rupture (LR=ΔF/Δt) (upper curve). The contact time refers to the time when the tip and surface are in constant contact (middle curve). The lower curve shows no binding event. (C) Box plot of specific binding probabilities measured by AFM between virions and JAM-A before and after injection of 10 μg/ml JAM-A antibody (AB). The horizontal line within the box indicates the median, boundaries of the box indicate the 25^th^ and 75^th^ percentile, and the whiskers indicate the highest and lowest values of the results. The square in the box indicates the mean. ns, *P* > 0.05; ****, *P* < 0.0001; determined by two-sample *t*-test in Origin. (D-F) Dynamic force spectroscopy (DFS) plot showing the distribution of rupture forces measured between a JAM-A-coated surface and rsT1L (D, black), Head (E, red), or HeadΔSA (F, light red), with average rupture forces determined for eight distinct LR ranges. Data corresponding to single interactions are fitted with the Bell-Evans (BE) model describing a ligand-receptor bond as a simple two-state model (I, black curve), providing average koff and xu values. Dashed lines represent the predicted binding forces for two (II) and three (III) simultaneous uncorrelated interactions (Williams-Evans model [WEM]). (G-I) The binding probability is plotted as a function of contact time. The line is the result of a least-squares fit of a monoexponential decay, providing average values for the kinetic on-rate of the probed interaction (k_on_). KD is calculated via k_off_/k_on_. Error bars indicate standard deviation of the mean. For all experiments, data are representative of *n* = 3 independent experiments.

### The Head mutant uncoats and transcribes viral positive-sense RNA with kinetics comparable to parental rsT1L and is sensitive to ammonium chloride

It is possible that crosslinking σ1 primes particles for disassembly and promotes conversion to the ISVP form. To test this hypothesis, we sought to quantify reovirus disassembly, transcription, and ammonium chloride (NH_4_Cl) sensitivity. When we initiated studies of in-cell reovirus disassembly, we determined that the Head mutant bound L cells poorly compared to rsT1L in our assay. This difference was surprising, given the enhanced avidity of the Head mutant for JAM-A (Fig. 6), but it may result from reduced SA-binding avidity (Fig. 4C), as initial σ1 binding to SA favors strong reovirus anchorage to JAM-A (18). To quantify the binding difference, we adsorbed L cells with rsT1L or 1, 5, or 25 times the number of Head mutant particles at 4°C for 1 h with rotation. After washing away unbound virus, we pelleted and lysed the cells and quantified bound virus by immunoblotting (Fig. 7A-B). Approximately 9.5 times as many Head mutant particles as rsT1L particles were required to achieve equivalent bound virus signal. Reduced binding was unexpected, given the enhanced replication kinetics and large-plaque phenotype of the Head mutant virus (Fig. 3).

**Figure 7.**
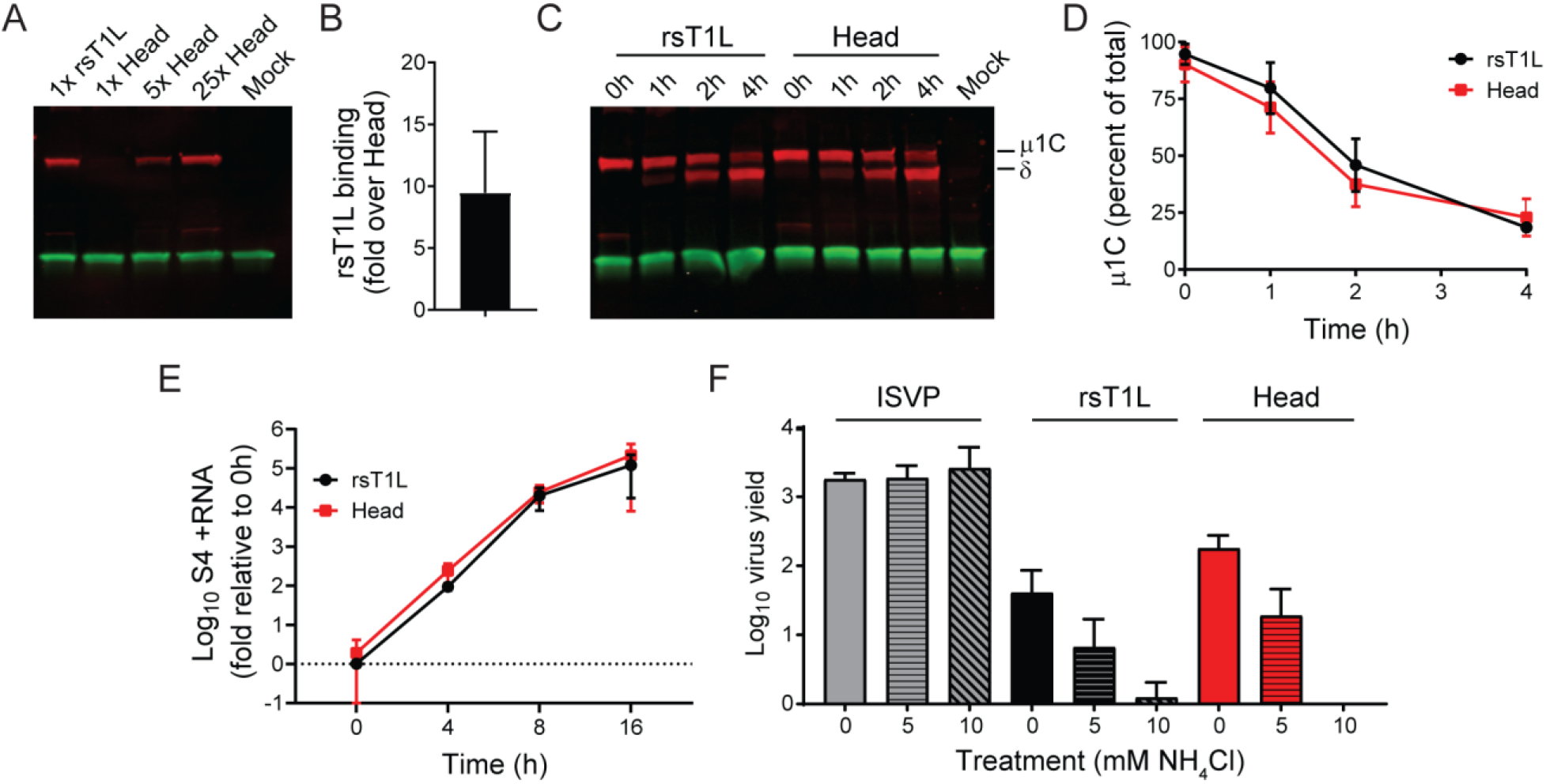
Head mutant uncoating and ammonium chloride sensitivity. (A) A fixed number of rsT1L viral particles and increasing ratios of Head mutant viral particles were adsorbed to L cells at 4°C for 1 h to allow cell binding but not entry. Following washing to remove unbound virus, cells were pelleted, and bound virus (red) was quantified using a LI-COR Odyssey following SDS-PAGE and immunoblotting using polyclonal reovirus antiserum and fluorescently labeled secondary antibody. pCNA antibody and fluorescently labeled secondary antibody were used to detect input cell lysate (green). (B) The mean and standard deviation of three independent experiments, as in (A), are shown. (C-D) Particle numbers of rsT1L and the Head mutant normalized for binding were incubated with L929 cells at 4°C for 1 h. Unbound virus was removed, and cells were incubated at 37°C to synchronize cell entry. After incubation for the times shown, cells were pelleted, and viral disassembly products μ1C and δ were quantified by immunoblotting as in (A). Shown are a representative immunoblot (C) and the mean and standard deviation of percent of μ1C at each time point from three quantified immunoblots (D). (E) L-cell monolayers were adsorbed with 100 PFU of virus particles at 4°C for 1 h to allow binding but not entry. Unbound virus was removed, and cells were incubated at 37°C with pre-warmed medium for 0, 4, 8, or 16 h prior to RNA extraction and purification and RT-qPCR analysis. Shown is the fold expression change of S4 +RNA relative to 0 h for the Head mutant and rsT1L, normalized to GAPDH. Shown are the mean and standard deviation from three independently infected wells per virus per time point analyzed in each of two independent experiments. (F) L cells were incubated in the presence of the concentrations of NH_4_Cl shown for 4 h prior to and following a 1 h adsorption with rsT1L ISVPs, rsT1L virions, or Head mutant virions at an MOI of 0.1 PFU/cell and a wash to remove unbound virus. At 24 h post-adsorption, cells were lysed by two rounds of freezing and thawing, and virus titers were determined by plaque assay. Shown are the mean and standard deviation from two virus clones per condition quantified in each of two independent experiments.

To identify differences in uncoating kinetics between rsT1L and the Head mutant, we adsorbed L cells with rsT1L or Head mutant particles yielding equivalent cell binding at 4°C for 1 h to permit binding but not entry. After washing away unbound virus, we added warm medium to synchronize entry, lysed cells during a time course, and quantified μ1 disassembly products μ1C and δ by immunoblotting (Fig. 7C-D). The kinetics of μ1C loss by rsT1L and the Head mutant were indistinguishable, suggesting that the presence of disulfide-mediated crosslinks in the σ1 Head domain does not alter uncoating kinetics. To identify differences in the transcription of viral positive-sense (+) RNA between rsT1L and the Head mutant, we used RTqPCR. We adsorbed L cells at 4°C with 100 PFU/cell of rsT1L or the Head mutant prior to wash and addition of pre-warmed medium, to allow synchronized cell entry. We incubated cells for 0, 4, 8, or 16 h at 37°C, harvested total RNA, and quantified reovirus S4 +RNA, normalized to GAPDH (Fig. 7E). Consistent with uncoating kinetics, S4 +RNA levels relative to those at 0 h were comparable between rsT1L and the Head mutant at every time point. Thus, σ1 crosslinks in the head domain fail to alter the kinetics of reovirus transcription.

Virions display reduced infectivity in the presence of NH_4_Cl, which prevents endosomal acidification, whereas ISVPs can directly penetrate cell membranes and are insensitive to the inhibitory effects of this compound (35, 36). To determine whether the Head mutant is sensitive to NH_4_Cl, L cells were incubated in the presence of 0, 5, or 10 mM NH_4_Cl for 4 h prior to and following adsorption with rsT1L ISVPs, rsT1L virions, or Head mutant virions. Unbound virus was removed by washing, cells were incubated for 24 h, and virus yields were quantified by plaque assay. While yields were slightly higher for the Head mutant in the absence of NH_4_Cl, both rsT1L and the Head mutant exhibited dose-dependent NH_4_Cl sensitivity (Fig. 7F). In contrast, ISVPs displayed no decrease in yield in the presence of NH_4_Cl. Together, these observations suggest that the enhanced replication kinetics and large-plaque phenotype of the Head mutant do not result from induction of an ISVP-like state through σ1 crosslinking.

## DISCUSSION

In this study, we discovered that restricting the conformational mobility of reovirus attachment protein σ1 by strategically introducing cysteine residues that crosslink the tail, body, or head domain (Fig. 2B) leads to either undetectable effects in L cells or an enhancement of both single-cycle replication kinetics and virus spread during multiple replication cycles, as indicated by plaque size (Fig. 3). These effects appear to be mediated not by alterations in σ1 encapsidation or gross misfolding (Fig. 2C-D), nor by particle aggregation (Fig. 4). For mutants with engineered cysteines in the head domain, JAM-A avidity is enhanced, independent of SA-binding avidity (Fig. 6). Head outer-capsid disassembly and viral +RNA transcription kinetics are identical to those of parental rsT1L (Fig. 7D-E), and particles require endosomal acidification to mediate infection (Fig. 7E). Together, these findings suggest that σ1 conformational changes are important mediators of reovirus attachment but do not influence the conversion from inactive to transcriptionally active particles during infection.

Our findings suggest that rsT1L σ1 conformational changes modulate reovirus attachment, but the mechanisms by which conformational changes are induced during infection remain unclear. Conversion of reovirus virions to ISVPs is associated with a conformational change in the σ1 protein (3, 20). Enhancement of JAM-A avidity has been reported for type 3 reovirus ISVPs and for type 3 reovirus virions following binding to SA, leading to the hypothesis that SA binding triggers the σ1 conformational changes observed in ISVPs (18). While it is unclear precisely why cysteine crosslinking of the rsT1L head domain enhances JAM-A avidity, a reasonable hypothesis is that the σ1 conformation induced by the formation of disulfide bridges in the head domain is ISVP-like, even though the rest of the particle resembles an intact virion. However, the trigger for this conformational change during infection of cells remains to be determined. T3D SA-binding and JAM-A-binding sites reside in the body and head domains, respectively, providing a potential mechanism for induction of conformational changes via “zipping up” of body or tail structures after SA binding (5, 18, 37). In contrast, T1L SA-binding and JAM-A-binding sites both reside in the head domain (4, 16), and mutations that significantly reduce SA binding fail to alter JAM-A avidity for HeadΔSA (Fig. 6). Thus, the trigger for T1L σ1 conformational changes is unclear. It is possible that yet unidentified cellular ligands bind rsT1L σ1 in the body or tail domain. Additionally, if σ1 undergoes a series of conformational changes during attachment and entry, crosslinking the head domain may induce a single step in a conformational cascade with multiple triggers, such as receptor binding, low pH, and proteolytic disassembly of the outer capsid.

Hemagglutination assays suggest that the Head mutant has decreased SA avidity (Fig. 4C), while AFM studies indicate it has enhanced JAM-A avidity (Fig. 6). Since the Head mutant was purified in the presence of NA, it is unclear whether the observed decrease in SA avidity results from crosslinking of the head domain, occupation of SA-binding sites, or both factors. In the case of HeadΔSA, which does not require NA for purification, diminished SA avidity in hemagglutination assays relative to rsT1L can be attributed to the engineered mutations in the SA-binding pocket (Fig. 4C). Despite its enhanced JAM-A avidity, the Head mutant binds poorly to cells relative to rsT1L (Fig. 7A-B). These observations suggest that efficient glycan binding contributes significantly to establishment of reovirus binding interactions with JAM-A at the cell surface. Reasons underlying the requirement for incubation with NA during Head mutant purification remain unclear. However, since HeadΔSA has a large-plaque phenotype, SA binding does not appear to be required to mediate enhanced spread in cultured cells (Fig. 4E).

Cysteine mutants that exhibit enhanced virus yields during a single round of replication also form the largest plaques (Fig. 3). Since all cysteine mutations promote disulfide bridge formation (Fig. 2B), and some mutations or pairs are in the same domain or even directly adjacent (Fig. 1B), it is unclear why only some of the mutations alter viral replication and plaque-size phenotypes (Fig. 3). However, differences in the efficiency of crosslinking may provide clues about σ1 conformation. Additionally, future studies aimed at determining the structure of the compact, virion-associated conformer of σ1 and the steps that mediate conformational changes during attachment and cell entry may help answer these questions.

What do observations to date suggest about the compact, virion conformation of σ1? Detection of disulfide bridges formed between the tail, body, and head domains of σ1 monomers suggests that the residues selected for cysteine mutagenesis are structurally adjacent to one another in purified reovirus particles under non-reducing conditions (Fig. 2B). This observation suggests that several tertiary and quaternary structures resolved by X-ray crystallography for fragments of purified, recombinant σ1 are maintained in its compact, virion form, even though the full-length model of σ1 based on these structures is thought to represent its ISVP form (Fig. 1) (8). For two cysteine mutants, Tail1 and Body1, disulfide bridges appear to be retained as dimers and trimers, respectively, even under reducing conditions (Fig. 2B). Asn38, which is exchanged with cysteine in Tail1, faces the interior of an α-helical coiled-coil close to the likely σ1 insertion point in the particle (6, 8). In this position, the Cys38 disulfide bridge may be protected from disruption, even in a reducing environment. For Body1, reasons for the relative stability of a disulfide bridge within the triple β-spiral region of σ1 are unclear but suggest that residues Gly287 and Val299 are protected in the virion conformation of σ1. The head domain appears to form monomers, dimers, and trimers under non-reducing conditions (Fig. 2B), which supports partial detrimerization or “breathing” of monomers within this region, as suggested previously in studies of a conserved aspartic acid motif in the head trimer interior (38). Structures of JAM-A in complex with σ1 suggest that the receptor engages a single monomer (16, 37). Yet, AFM data clearly indicate that JAM-A avidity is higher for the Head mutant than for rsT1L (Fig. 6). These data suggest that the JAM-A-binding surface of σ1 in the virion is not highly accessible but becomes more accessible in the ISVP or the Head mutant. Together, these observations support a model in which the σ1 protein is at least partially detrimerized on the virion, perhaps with the head domains facing towards the particle surface. Some event, such as binding to SA or an unidentified receptor, may induce conformational changes that promote σ1 extension and trimerization of the head domain, thereby enhancing accessibility of the JAM-A binding site. Reductions in pH in the endocytic compartment also have been hypothesized to promote trimerization of the σ1 head domain during entry and may function as a trigger of conformational change (38).

The capacity of σ1 to dissociate into a monomeric form does not appear to be required for reovirus replication. In fact, there may be a replication advantage to maintaining a dimeric or trimeric conformation. So, why have reoviruses not evolved to do so? One possibility is that multimerized forms of σ1 have less avidity for SA, which is suggested by the reduced capacity of Head to agglutinate red blood cells (Fig. 4C). Reduced SA-binding capacity may impair overall binding of viruses to cells by preventing the initial low-affinity interactions that permit lateral diffusion (17). Additionally, although Tail1, Body2, and Head exhibit enhanced replication kinetics and form larger plaques in L cells, relative to rsT1L (Fig. 3), it is unclear whether these viruses would maintain an advantage *in vivo.*

Many viruses undergo stepwise conformational rearrangements during cell attachment and entry that can be triggered by cellular factors such as receptors, proteolytic enzymes, or pH (39, 40). In many cases, these rearrangements facilitate attachment, penetration of a cell membrane, such as the endosomal membrane, or release of the viral genome from the particle. For example, the rotavirus spike attachment protein, VP4, is activated by proteolytic cleavage, which triggers a distinct conformational reorganization (41, 42). Proteolysis is thought to prime VP4 for an even more dramatic conformational rearrangement resembling that of enveloped virus fusion proteins, which mediates endosomal penetration and is induced during uncoating. The current study underscores the functional significance of viral structural protein dynamics. It is likely that many viruses employ conformational rearrangements of outer-capsid proteins, including attachment proteins, to modulate early replication processes.

## MATERIALS AND METHODS

### Cells

Spinner-adapted L cells were grown in suspension culture in Joklik’s minimum essential medium (US Biological) supplemented to contain 5% fetal bovine serum (FBS) (Gibco), 2 mM L-glutamine (Corning), 100 units/ml penicillin/100 μg/ml streptomycin (Corning), and 25 ng/ml amphotericin B (Corning). Baby hamster kidney cells expressing T7 RNA polymerase under control of a cytomegalovirus promoter (BHK-T7) (43) were maintained in Dulbecco’s minimum essential medium (Corning) supplemented to contain 5% FBS (Gibco), 2 mM L-glutamine (Corning), 100 units/ml penicillin/100 μg/ml streptomycin (Corning), and 25 ng/ml amphotericin B (Corning), with 1 mg/ml Geneticin (Gibco) added during alternate passages.

### Viruses

Laboratory stocks of parental reovirus strain rsT1L and σ1 cysteine mutants were engineered using plasmid-based reverse genetics (43, 44). Sites for introduction of cysteine pairs in body and head domains were selected using Disulfide by Design 2 (45). The pBacT7-S1T1L plasmid was used as a template to engineer pBacT7-S1T1L N38C (Tail1), pBacT7-S1T1L N94C (Tail2), pBacT7-S1T1L G287C, V299C (Body1), pBacT7-S1T1L D89C, S301C (Body2), pBacT7-S1T1L M332C, S403C (Head), and pBacT7-S1T1L M332C, S370P, Q371E, S403C (HeadΔSA) by ‘round the horn PCR (46) with mutagenic primers. Monolayers of ∼ 8 × 10^5^ BHK-T7 cells were co-transfected using TransIT-LT1 transfection reagent (Mirus Bio LLC) with 0.8 μg each of nine plasmid constructs representing the T1L reovirus genome plus a single parental or mutant pBacT7-S1 plasmid. After a minimum of five days of incubation, recombinant viruses were isolated by plaque purification on L cells. Virus stocks were prepared from at least two plaque-purified clones per recombinant virus strain (47). ISVPs were prepared by treatment of virions with chymotrypsin (Sigma) (48). Viral titers were determined by plaque assay using L cells, as described (47). Chymotrypsin (Worthington Biochemical, 10 μg/ml) was added to plaque assay overlays for some experiments.

### Reovirus purification

Virus particles were purified from infected L cells by deoxycholate permeabilization, Vertrel XF (DuPont) extraction, and CsCl gradient centrifugation, as described (20). Reovirus particle concentration was determined from the equivalence of one unit of optical density at 260 nm to 2.1 × 10^12^ particles. To purify the Head mutant, 14 mU of *Arthrobacter ureafaciens* NA (Roche) was incubated with the infected cell pellet at room temperature for 1 h prior to treatment with deoxycholate.

### Immunoblot analysis of σ1 multimeric status

Purified reovirus particles (2 x 10^10^) were boiled in sample buffer containing (reducing) or lacking (non-reducing) 100 mM β-mercaptoethanol and resolved by SDS-PAGE in a 4-20% Mini-PROTEAN TGX Precast Protein Gel (BIO-RAD). Proteins were transferred to nitrocellulose membranes using a Trans-Blot SD Semi-Dry Transfer Cell (BIO-RAD) at 20 V for 30 min. Membranes were blocked in Odyssey blocking buffer and incubated with rabbit anti-T1L σ1 head serum diluted 1:500 (49). Membranes were washed in PBS-T (PBS + 0.1% Tween-20) and incubated with anti-rabbit IRDye680LT Abs (LI-COR) diluted 1:15,000 in PBS-T + 5% non-fat dry milk. After washing with PBS-T, membranes were scanned using an Odyssey infrared imaging system (LI-COR).

### Agarose gel separation of intact virions

Purified reovirus virions (2.5 × 10^11^) were diluted into TAE buffer (40 mM Tris acetate, 1 mM EDTA [pH 8.0]) containing 2.5% glycerol and 0.025% bromophenol blue. Particles were resolved by electrophoresis in 1% agarose in TAE buffer at room temperature for 18 h (50). Bands representing reovirus particles containing 0-12 σ1 trimers were visualized using PageBlue Protein Staining Solution (Thermo Scientific).

### Assessment of reovirus infectivity by fluorescence imaging

L cells were seeded into 96-well, black-walled plates to achieve a density of ∼ 2 × 10^4^ per well and adsorbed with serial dilutions of virus at 37°C for 1 h. Inocula were removed, and cells were washed and incubated in fresh medium at 37°C for 16-20 h. Cells were fixed with cold methanol, and reovirus proteins were detected by incubation with polyclonal reovirus antiserum at a 1:500 dilution in PBS containing 0.5% Triton X-100 at 37°C, followed by incubation with Alexa Fluor 488-labeled secondary IgG (Invitrogen) and DAPI. Images were captured for four fields of view per well using an ImageXpress Micro XL automated microscope imager (Molecular Devices). Total and percent infected cells were quantified using MetaXpress high-content image acquisition and analysis software (Molecular Devices). The effect of σ1 blockade (microneutralization) was tested by incubating virions at particle concentrations that yielded approximately 80% infected cells per well with medium or medium containing the indicated concentrations of mAb 5C6 (27, 51), donkey anti-mouse IgG (Jackson Immuno Research), or polyclonal reovirus antiserum at 37°C for 1 h prior to adsorption onto L cells, incubation, fixation, staining, and quantification.

### Quantifying virus replication

L cells were seeded into wells of 12-well plates to achieve a density of ∼ 8 × 10^5^ per well and adsorbed with reovirus in triplicate at an MOI of 0.1 PFU/cell at room temperature for 1 h. Inocula were removed, and cells were incubated in fresh medium at 37°C for 0, 4, 8, 12, 16, 20, or 24 h prior to two rounds of freezing and thawing. Viral titers in inocula and cell lysates were determined by plaque assay in L cells.

### Dynamic light scattering

rsT1L or cysteine mutants (2 × 10^12^ particles/ml) were analyzed using a DynaPro NanoStar (Wyatt Technology). Measurements were made at room temperature using quartz cuvettes. Hydrodynamic radii were determined by averaging readings from 3 × 10 iterations per sample, each with a 5 s acquisition time, using DYNAMICS software (Wyatt Technology). No filters were applied to the data.

### Cell-binding, in-cell uncoating assay

To quantify cell binding, L cells (10^6^) were aliquoted into Eppendorf tubes on ice. Virus dilutions (10^4^ particles/cell) were prepared in cold medium. For the Head mutant, virus dilutions at 5 × 10^4^ and 2.5 × 10^5^ particles/cell also were prepared. Cells were pelleted, adsorbed with virus at 4°C for 1 h, pelleted a second time, and washed twice prior to the addition of RIPA buffer supplemented with 1 mM PMSF. Samples were boiled with SDS sample buffer and resolved on a 10% Mini-PROTEAN TGX Precast Protein Gel (BIO-RAD). Proteins were transferred to nitrocellulose membranes, which were blocked in Odyssey blocking buffer, then incubated with 1:2000 diluted mouse monoclonal anti-PCNA (Santa Cruz, ∼34 kDa) and 1:1000 diluted rabbit polyclonal anti-reovirus serum. Membranes were washed in PBS-T and incubated with anti-rabbit IRDye680LT Abs and anti-mouse IRDye800CW Abs (LI-COR) diluted 1:15,000 in PBS-T + 5% non-fat dry milk. After washing with PBS-T, membranes were scanned using an Odyssey infrared imaging system (LI-COR). Protein band intensity was quantified using Image Studio version 5.2 in three independent experiments.

To quantify reovirus in-cell uncoating, L cells (10^6^) were aliquoted into Eppendorf tubes on ice. Virus dilutions of 10^4^ rsT1L particles/cell and Head mutant particles resulting in equivalent cell binding were prepared in cold medium. Cells were pelleted, adsorbed with virus at 4°C for 1 h, pelleted a second time, and washed twice prior to addition of pre-warmed medium, to allow synchronized cell entry. Cells were incubated for 0, 1, 2, or 4 h before pelleting and resuspension in RIPA buffer supplemented with 1 mM PMSF. Samples were resolved and proteins were detected and quantified as in the cell-binding assay. Integrated protein band fluorescence for μ1C was divided by that for total μ1 to quantify uncoating in three independent experiments.

### Ammonium chloride sensitivity

To quantify reovirus replication in the presence of ammonium chloride, L cells (5 × 10^5^) were pre-incubated in medium containing 0, 5, or 10 mM NH_4_Cl at 37°C for 4 h prior to adsorption with 0.1 PFU/cell of rsT1L, Head, or HeadΔSA virions or rsT1L ISVPs at room temperature for 1 h. Cells were washed and incubated in medium containing 0, 5, or 10 mM NH_4_Cl at 37°C for 0 or 24 h. Cells were lysed by two rounds of freezing and thawing, and viral titers in cell lysates were determined by plaque assay in L cells.

### Functionalization of AFM tips

AFM tips (MSCT probes, Bruker) were coated with virions using flexible PEG linkers (NHS-PEG27-aldehyde) in a three-step coupling procedure as described (52). Amino-functionalization of the tips (step 1) was conducted using aminopropyltriethoxysilane (APTES) coupling in gas phase (53). First, the cantilevers were rinsed in chloroform (Sigma-Aldrich) for 5 min and cleaned in UV radiation and ozone (UV-O) (Jetlight) for 15 min. The AFM tips were placed in a desiccator under argon atmosphere together with trays containing 30 μl APTES and 10 μl of trimethylamine, respectively, for 2 h. After removal of the APTES and triethylamine trays, the AFM cantilevers were left to cure under argon for at least 2 days. After amino-functionalization, the NHS-PEG27-aldehyde linker was coupled to the AFM tip as follows (step 2). The cantilevers were immersed for 2 h in a solution prepared by mixing 3.3 mg of NHS-PEG27-aldehyde linker dissolved in 0.5 ml of chloroform with 30 μl of triethylamine, washed with chloroform, and dried with a stream of filtered nitrogen gas. In step 3, the tips were placed on Parafilm in a polystyrene Petri dish, stored within an icebox, followed by pipetting of 100 μl virus solution (10^9^ particles/ml) onto the cantilevers and adding 2 μl of a freshly prepared sodium cyanoborohydride solution (∼ 6% wt/vol in 0.1 M NaOH_(aq)_) to the virus droplet. The tips were incubated in 4°C for 1 h. Ethanolamine (5 μl, 1 M, pH 8.0) was added and mixed carefully with the drop on the cantilevers, followed by incubation at 4°C for 10 min to quench the reaction. Subsequently, tips were washed three times in ice-cold virus buffer (150 mM NaCl, 15 mM MgCl_2_, 10 mM Tris, pH 7.4) and stored in individual wells of a multiwell dish containing 2 ml of ice-cold virus buffer per well.

### Preparation of JAM-A-coated model surfaces

His_6_-tagged JAM-A (Bio-Connect Life Science) was immobilized using NTA-His_6_ binding chemistry. The gold-coated surfaces were rinsed with ethanol, dried with a gentle nitrogen flow, and cleaned using a 15 min UV radiation and ozone (UV-O) treatment (Jetlight). Surfaces were immersed overnight in an ethanol solution containing 0.05 mM of NTA-terminated (10%) and PEG-terminated (90%) alkanethiols. The next day, model surfaces were rinsed with ethanol and immersed for 1 h in 40 mM aqueous solution of NiSO4 (pH 7.2). After rinsing the surfaces with water, they were incubated with JAM-A protein (0.1 mg/ml) for 1 h. The sample surfaces were rinsed 10 times with PBS and used immediately or stored at 4°C, always keeping the surfaces hydrated.

### Force distance-based AFM on model surfaces

Force distance (FD) curve-based AFM experiments were conducted at room temperature using virus-functionalized MSCT-D probes (spring constants were calculated using thermal tune (54), with values ranging from 0.027 to 0.043/nm). Model surfaces, grafted with JAM-A as described (18), were mounted on a piezoelectric scanner using a magnetic carrier immersed in PBS. AFM Nanoscope Multimode 8 (Bruker, Nanoscope software v9.1) and Force-Robot300 (Bruker, Germany) were used to conduct these experiments. The FD curves were recorded from 5 × 5 μm arrays in the force-volume (contact) mode, with 32 x 32 pixels resolution (corresponding to 1024 FD curves). For all experiments, the ramp size was set to 500 nm, and the maximum force was limited to 500 pN, while the approach velocity was kept constant at 1 μm/s.

Dynamic force spectroscopy (DFS) experiments were conducted with no surface delay and, to probe a wide range of loading rates, the retraction velocities were varied and set to 0.1, 0.2, 1, 5, 10 and 20 μm/s. To ensure specific interactions between the functionalized tip and sample, an independent negative control was included by surface blocking: measurements were obtained before and after injection of a JAM-A antibody (Sigma Aldrich, concentration of 10 μg/ml) to the sample surface to block JAM-A binding. The same position on the surface was probed several times using the same tip with a retraction velocity of 1 μm/s. In a different set of experiments, measurements of the kinetic on-rate constant (*k_on_*) were conducted by allowing the cantilevers to rest on the surface for different intervals (starting with no pause time, data were collected with contact times of 50 ms, 100 ms, 150 ms, 250 ms, 500 ms, and 1 s).

### Data analysis

Depending on the instrument used, either JPK Data Processing (version 6.1.149) or Nanoscope software (version 9.1) was employed for analysis. To identify peaks corresponding to adhesion events occurring between particles linked to the PEG spacer and the JAM-A model surface, the retraction curve before bond rupture was fitted with the worm-like chain model for polymer extension (55). Regarding DFS experiments, data including loading rates and disruption forces were extracted. Origin software (OriginLab) was used to display the results in DFS plots to fit histograms of rupture force distributions for distinct loading rate ranges and to apply various force spectroscopy models as described (32, 33). For kinetic on-rate analysis, the binding probability (BP) (fraction of curves showing binding events) was determined at a certain dwell time (*t*) (time the tip is in contact with the surface). Those data were fitted and K_D_ calculated as described previously (56, 57).

### Quantitative reverse transcription PCR

To quantify reovirus S4 +RNA during a time course, L cells (2 × 10^6^) were chilled and adsorbed with 100 PFU/cell of rsT1L or Head mutant virus at 4°C for 1 h. Then, inocula were aspirated, and cells were washed prior to addition of pre-warmed medium, to allow synchronized cell entry. Cells were incubated at 37°C for 0, 4, 8, or 16 h before RNA was harvested by direct cell lysis in TRIzol reagent (Invitrogen) and RNA purification using the PureLink RNA Mini Kit (Invitrogen), according to the manufacturer’s protocol. Total RNA was quantified using a Qubit fluorometer (Invitrogen). Then, reovirus S4 +RNA in 500 ng of total RNA was quantified using the Superscript III First Strand cDNA Synthesis System (Invitrogen) and PowerUp SYBR Green Master Mix (Applied Biosystems), according to manufacturer protocols with minor modification. Each reaction was performed in duplicate. To quantify S4 +RNA, we used an S4 reverse primer in the reverse transcription reaction and S4-specific forward and reverse primers (58) in the qPCR reaction. To quantify GAPDH for sample normalization, we used oligo(dT) to amplify cellular mRNAs in the reverse transcription reaction and GAPDH-specific forward and reverse primers (59) in the qPCR reaction. Primers were annealed at 65°C for 5 min then cooled to 4°C. Reverse transcription was performed with aby incubation at 25°C for 5 min then at 55°C for 45 min. The reaction was terminated by incubation at 70°C for 15 min. Subsequently, 40 cycles of quantitative PCR were performed at 95°C for 15 s followed by incubation at 60°C for 1 min using a StepOnePlus Real-Time PCR System (Applied Biosciences). Product specificity was checked by dissociation curve analysis. The relative quantity of reovirus S4 +RNA was determined. We calculated *ΔΔC_T_* relative to rsT1L at each time point as (unknown *C_Ttx_* - GAPDH *C_Ttx_*) - (*rsT1L C_Ttx_* - GAPDH *C_Ttx_*). We calculated *ΔΔC_T_* for each sample as (unknown *C_Ttx_* - GAPDH *C_Ttx_*) - (unknown *C_Tt0_* - GAPDH *C_Tt0_*), where *C_T_* is the threshold cycle, *t0* is the time of infection, and *tx* is the time post-infection. The relative quantity of reovirus S4 +RNA was determined by raising 2 to the negative logarithm of *ΔΔC_T_*.

### Hemagglutination assay

Purified virions were aliquoted into a 96-well U-bottom microtiter plate (Costar), starting with a concentration of 1.25 × 10^10^ particles/well, and serially diluted 1:2 in 50 μl PBS. Type O human erythrocytes were washed twice with PBS, resuspended to 1% (v/v) in PBS, added to virus-containing wells at 50 μl/well, and incubated at 4°C for 4 h. One HA unit is the minimum particle number that produces a partial or complete erythrocyte shield in the well. HA titer is defined as 2.5 × 10^10^ particles divided by the number of particles per HA unit.

### Statistical analysis

Statistical analyses were conducted using Prism 8.2.1 (GraphPad). Plaque sizes, μ1C accumulation (uncoating), S4 positive-sense RNA transcription, and virus yield during a time course were compared between reovirus mutants and rsT1L using an unpaired Student’s *t* test. For uncoating, S4 positive-sense RNA transcription, and virus yield, discovery was determined using the two-stage linear step-up procedure of Benjamini, Krieger, and Yekutieli, with a false discovery rate (Q) of 1%. Each time point was analyzed individually, without assuming a consistent standard deviation.

## ACKNOWLEDGEMENTS

We thank James M. Hutchison from the Charles R. Sanders lab in the Department of Biochemistry at Vanderbilt University School of Medicine for assistance with dynamic light scattering experiments. We thank members of the Dermody, Ogden, and Stehle laboratories for many helpful discussions during the conduct of this work. We are grateful to Danica M. Sutherland and Gwen M. Taylor for careful review of the manuscript.

This work was supported by U.S. Public Health Service award R01 AI118887 to Terence S. Dermody, Kristen M. Ogden, and Thilo Stehle. Work at the Université Catholique de Louvain was supported by the European Research Council (ERC) under the European Union’s Horizon 2020 research and innovation programme (Grant Agreement 758224), the FNRS-WELBIO (Grant WELBIO-CR-2019S-01), the National Fund for Scientific Research (FNRS), and the Research Department of the Communauté francş ise de Belgique (Concerted Research Action). D.A. is Research Associate at the FNRS. The funders had no role in study design, data collection and interpretation, or the decision to submit the work for publication.

